# The Golgi-associated retrograde protein (GARP) complex plays an essential role in the maintenance of the Golgi glycosylation machinery

**DOI:** 10.1101/2020.12.21.423858

**Authors:** Amrita Khakurel, Tetyana Kudlyk, Juan S. Bonifacino, Vladimir V. Lupashin

## Abstract

The Golgi apparatus is a central hub for intracellular protein trafficking and glycosylation. Steady-state localization of glycosylation enzymes is achieved by a combination of mechanisms involving retention and vesicle recycling, but the machinery governing these mechanisms is poorly understood. Herein we show that the Golgi-associated retrograde protein (GARP) complex is a critical component of this machinery. Using multiple human cell lines, we show that depletion of GARP subunits is detrimental to N- and O-glycosylation, and reduces the stability of glycoproteins and Golgi enzymes. Moreover, GARP-KO cells exhibit impaired retention of glycosylation enzymes in the Golgi. Indeed, a RUSH assay shows that, in GARP-KO cells, the enzyme beta-1,4-galactosyltransferase 1 is not retained at the Golgi but instead is missorted to the endolysosomal compartment. We propose that the endosomal compartment is part of the trafficking itinerary of Golgi enzymes and that the GARP complex is essential for recycling and stabilization of the Golgi glycosylation machinery.

## Introduction

The Golgi complex plays a central role in the processing, packaging and sorting of secretory and transmembrane proteins in eukaryotic cells (1) (2). Although the Golgi complex was discovered more than 120 years ago, the molecular mechanisms of intra-Golgi trafficking and Golgi maintenance are not entirely understood (2). According to the “cisternal maturation” model, newly-synthesized secretory proteins arrive at the *cis*-Golgi compartment from the ER by vesicular trafficking, and then stay within this compartment while it “matures”, sequentially changing its identity from *cis* to medial to *trans* (3) (4). During this maturation, Golgi-resident glycosylation enzymes are recycled continuously from late to early cisternae by retrograde trafficking, allowing them to maintain their steady-state localization in the Golgi (5) (6). However, the exact trafficking itineraries and molecular machinery involved in the delivery of these enzymes to their corresponding Golgi cisternae are poorly understood. Potential components of this machinery are coat proteins, small GTPases, vesicle tethers, and SNAREs implicated in inter-compartmental cargo transport in both anterograde and retrograde pathways (7) (1) (8). Each Golgi cisterna contains different carbohydrate-modifying enzymes. These enzymes catalyze the addition (glycosyltransferases) or removal (glycosidases) of sugars to/from cargo glycoproteins, as well as the addition of sulfate and phosphate groups. As a maturing glycoprotein proceeds from *cis*- to *trans*-Golgi, several Golgi glycosyltransferases add *N*-acetylglucosamine, galactose and sialic acid, resulting in proper glycosylation of the protein (9).

The conserved oligomeric Golgi (COG) complex is the major Golgi vesicle tethering complex responsible for the maintenance of the Golgi glycosylation machinery (1) (10). Several lines of evidence indicate that COG and another vesicle tethering complex, the Golgi-Associated Retrograde Protein (GARP) complex, are functionally interconnected (11) (12). In addition, multiple recent CRISPR screens identified both COG and GARP as a requirement for the entry of viruses and bacterial toxins into the host cells (13) (14) (11). These pathogens interact with glycoproteins or glycolipids on the host plasma membrane, suggesting that complete depletion of COG and GARP complexes may alter not only a particular intracellular trafficking step, but also the cellular glycosylation machinery.

GARP is a multi-subunit tethering complex (MTC) located at the *trans*-Golgi network (TGN) where it functions to tether retrograde transport vesicles derived from endosomes (15) (16) (17) (18) (19). GARP comprises four subunits named vacuolar protein sorting 51 (VPS51), VPS52, VPS53, and VPS54 (15) (16) (17) (18) (20). GARP shares its VPS51, VPS52 and VPS53 subunits with another complex known as Endosome-Associated Recycling Protein (EARP/GARPII) complex, which has an additional subunit named VPS50 in place of VPS54 (21) (22). GARP complex localization to the TGN is dependent on the small GTPases ARFRP1 and ARL5 (23) (24). In yeast, mutation of genes encoding any GARP subunit inhibits recycling of the cargo receptor VPS10 from endosomes to the TGN, leading to the missorting and secretion of vacuolar hydrolases (15) (16). In mammalian cells, GARP participates in retrieval of mannose-6-phosphate receptors (MPRs), the TGN-resident protein TGN46 and the Niemann-Pick C2 (NPC2) protein from endosomes to the TGN (18) (25). As a consequence of MPR missorting, GARP-deficient cells exhibit increased secretion of immature cathepsin D with subsequent lysosomal dysfunction (18). GARP depletion also results in alterations of autophagy (20) (26), anterograde transport of GPI-anchored and transmembrane proteins (27) and sphingolipid homeostasis (28). Furthermore, mutations in GARP subunits have been found to cause neurodevelopmental disorders in humans (29) (30) (31) (32) and a mutation in VPS54 is the cause of embryonic lethality with progressive motor neuron death in the wobbler mouse, an animal model for Amyotrophic Lateral Sclerosis (ALS) (33). Interestingly, inhibition of sphingolipid synthesis showed improvement in neuropathology and survival in mutant wobbler mice, suggesting that altered sphingolipid homeostasis underlies the pathogenesis of GARP-deficiency disorders (34).

In this study, we have examined if glycosylation is disturbed in human cells depleted of GARP subunits. Using three different cell lines with knock-out (KO) of GARP subunits and a combination of microscopy and biochemical approaches, we demonstrate defects in glycosylation and reduction in the level of Golgi-resident enzymes in GARP-KO cells. We also show that the *trans*-Golgi enzymes beta-1,4-galactosyltransferase 1 (B4GalT1) and alpha-2,6-sialyltransferase 1 (ST6Gal1) failed to be retained in the Golgi but are instead mislocalized to endolysosomes in GARP-KO cells. These findings indicate that GARP plays a crucial role in normal Golgi glycosylation by mediating the maintenance of the Golgi glycosylation machinery.

## Materials and Methods

### Cell culture

hTERT RPE1 (retinal pigment epithelial) and HEK293T cells used for all experiments were purchased from ATCC. HeLa-KO used in the study were previously described (24). RPE1, HEK293T and HeLa cells were cultured in Dulbecco’s Modified Eagle’s Medium (DMEM) containing Nutrient mixture F-12 (Corning) supplemented with 10% Fetal Bovine Serum (FBS) (Thermo Fisher). Cells were incubated in a 37°C incubator with 5% CO_2_ and 90% humidity.

**Table 1:**
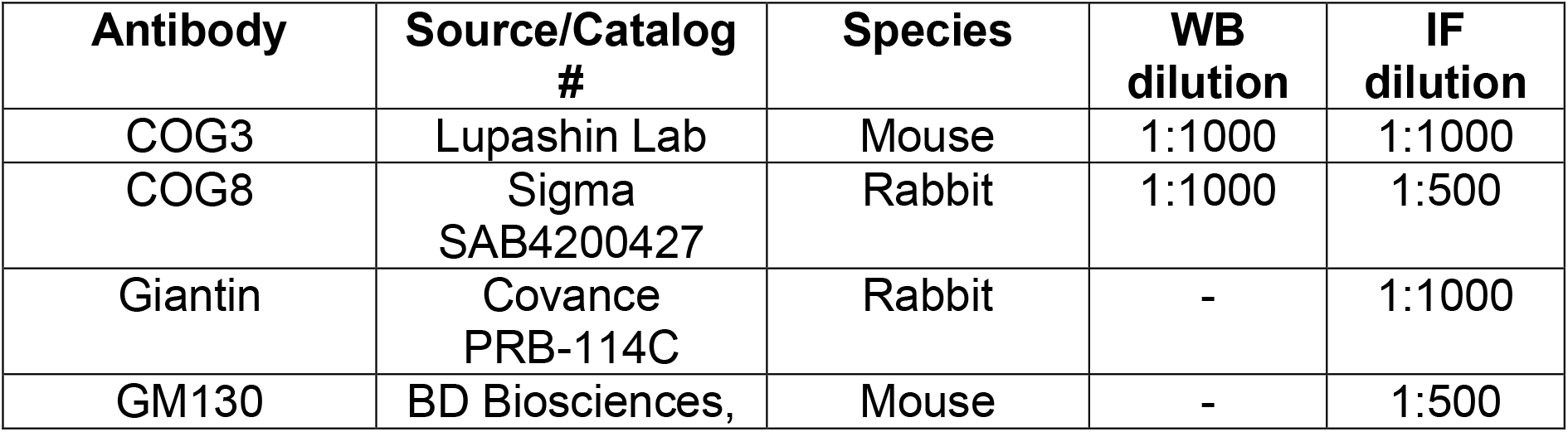
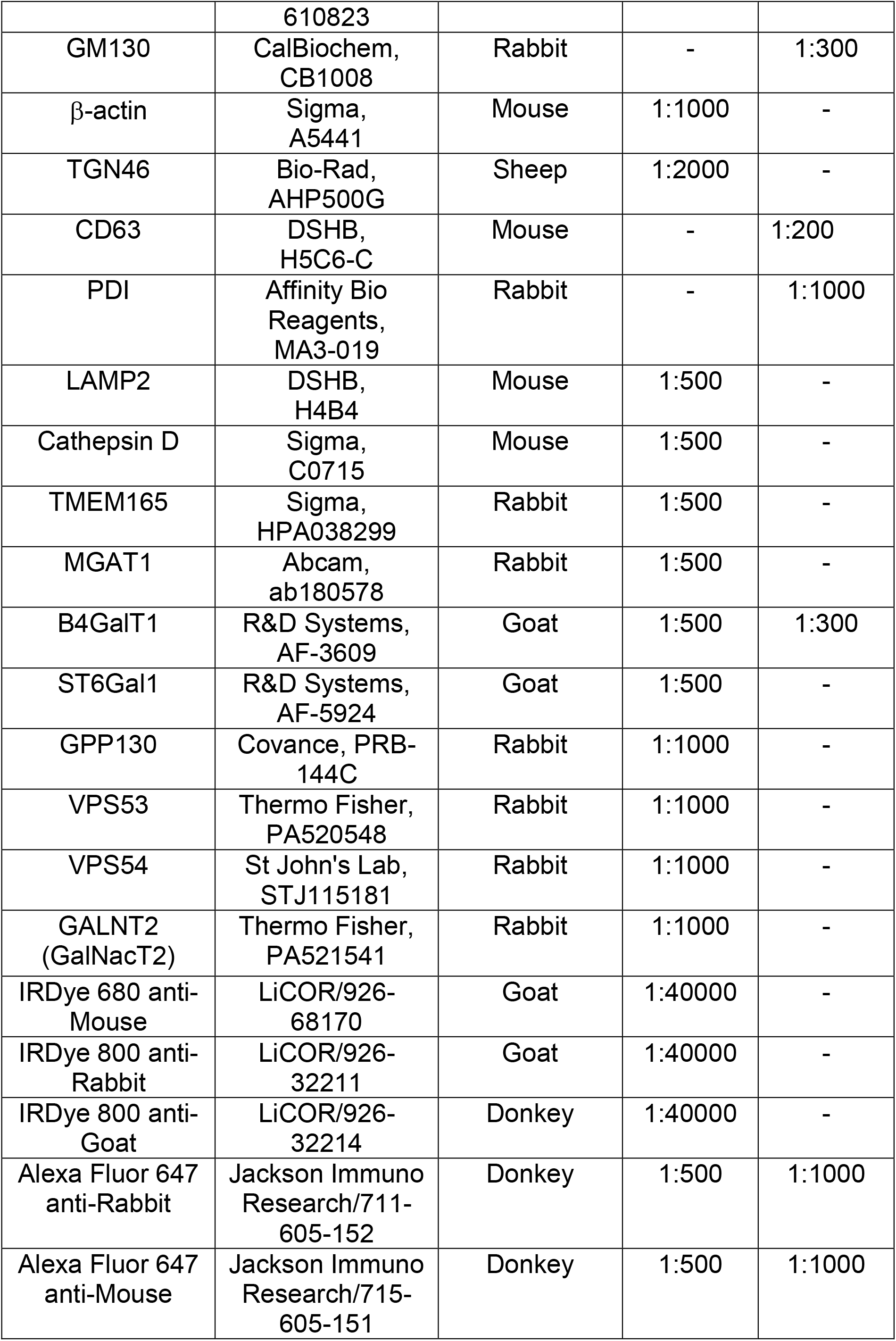
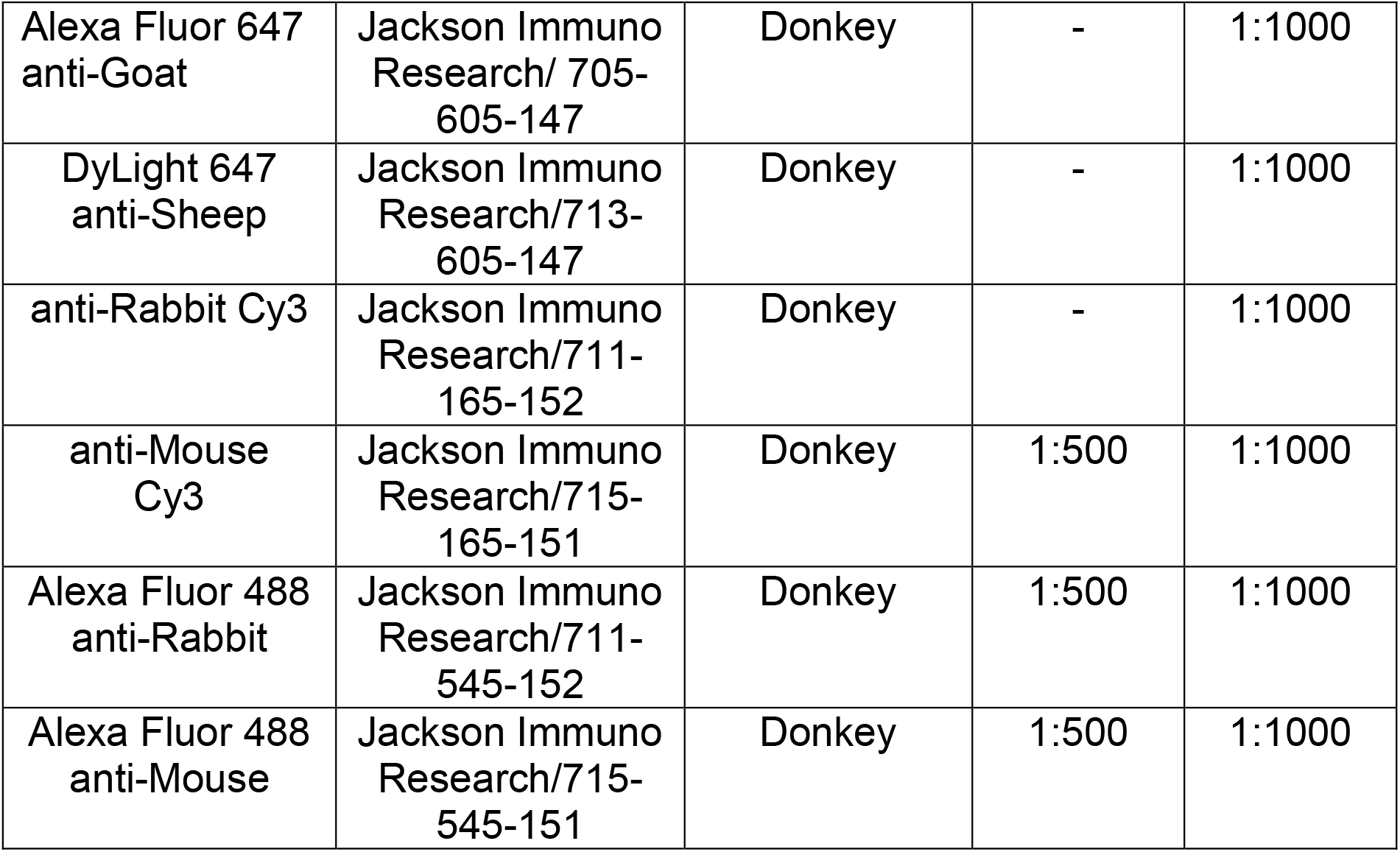
List of antibodies used.

**Table 2:** List of plasmids used.

| **Plasmid Name** | **Source** | **Citation** |
| --- | --- | --- |
| Lenti Cas-9 blast | Addgene #52962 Feng Zhang | Sanjana et al., 2014 (35) |
| hVPS53-GFP | Origene |  |
| pCl-neo-VPS54-13myc | Juan Bonifacino | Bonifacino et al., 2019 (24) |
| pENTR1A no ccDB (w48-1) | Eric Campeau and Paul Kaufman, Addgene #17398 | Campeau et al., 2009 (36) |
| pLenti CMV Neo DEST (705-1) | Eric Campeau and Paul Kaufman, Addgene #17392 | Campeau et al., 2009 (36) |
| pMD2.G | Didier Trono. Addgene #12259 | Dull et al., 1998 (37) |
| pRSV-Rev | Didier Trono. Addgene #12253 | Dull et al., 1998 (37) |
| pMDLg/pRRE | Didier Trono. Addgene #12251 | Dull et al., 1998 (37) |
| hVps53 - GFP pLenti COG4 promoter Neo DEST | This study | This study |
| mVps54 -13myc pLenti COG4 promoter Neo DEST | This study | This study |
| pLenti COG4 promoter Neo DEST (705-1) | This study | This study |
| LAMP2-GFP | Santiago Di Pietro | Ambrosio et al., 2012 (38) |
| ST-RFP | James Rothman | Lavieu et al., 2013 (39) |
| Str-KDEL_flgalT-SBP-mCherry | Franck Perez Addgene #65273 | Boncompain et al., 2012 (40) |
| mCherry-Acta | Adam D. Linstedt | Willett et al., 2013<br>(41) |

### Creation of VPS53- and VPS54-KO stable cell lines

To create RPE1 VPS53 and VPS54 stable knockouts, dual gRNAs were purchased from Transomic with the following target sequences:

### For VPS53-KO

transEDIT-dual CRISPR for VPS53 (TEDH-1088057)

grna-a: GCATTGAAATCTGCTCGATCTAGAGGGTCC

grna-b: TGACAATATTCGAACTGTTGTAAGAGGTCA

transEDIT-dual CRISPR for VPS53 (TEDH-1088055)

grna-a: ATGCACCACTCACGTGGAAACATGCGGCCG

grna-b: CTGGAGCACTTCCACAAGTATATGGGGATT

transEDIT-dual CRISPR for VPS53 (TEDH-1088056)

grna-a: CCAAGATTATGCGTACCAGAGCGAAGGAAA

grna-b: CCGCAGATCCGGCAGCTTTCCGAAAGGTAA

### For VPS54-KO

transEDIT-dual CRISPR for VPS54 (TEDHG1001)

grna-a: ACAAATATTCCTGAAACAGGCAGAAGGAAC

grna-b: ATCTAGAAAGTGTTATGAATTCCATGGAAT

transEDIT-dual CRISPR for VPS54 (TEDHG1001)

grna-a: CAAAAGATAATTCACTGGACACAGAGGTGG

grna-b: CATTCTACCTCCCACAGATCAGCAAGGAAC

transEDIT-dual CRISPR for VPS54 (TEDHG1001)

grna-a: CTTAACTCTGTAGCCACAGAAGAAAGGAAA

grna-b: GTAAGCATGTCAGTAGTAACAGATGGGATG

First, RPE1-Cas9 stable cell line was created by lentiviral transduction with a plasmid encoding FLAG-tagged Cas9. HEK293FT cells were used for the generation of lentiviral particles. Equal amounts of the three lentiviral packaging plasmids pMD2.G, pRSV-Rev, and pMDLg/pRRE, plus destination plasmid pLenti Cas9-Blast, were used. Briefly, transfected HEK293FT cells were placed in serum-reduced Opti-MEM with 25 μM Chloroquine and GlutaMAX. The next day, the medium was changed to Opti-MEM supplemented with GlutaMAX. At 72 h after transfection, the medium was collected, and cell debris removed by centrifugation at 600×g for 10 min. The supernatant was passed through 0.45 μM polyethersulfone (PES) membrane filter and lentiviral medium was stored at 4°C overnight. 1 ml of this lentiviral medium was used to transduce RPE1 cells on a 10 cm dish along with 50 mM sodium butyrate. After 24 h, the medium was changed to DMEM/F12 supplemented with 10% FBS and 10 μg/ml blasticidin.

After the creation of hTERT RPE-Cas9 stable cell line, these cells were transfected with a cocktail of three dual gRNAs specific for VPS53 and VPS54. Neon electroporation (Thermo Fisher) was used for transfecting cells according to the manufacturer’s protocol. TransEDIT-dual CRISPR plasmids express GFP; therefore, the efficiency of transfection was estimated by counting GFP-positive cells. At 48 h post-transfection, cells were spun down at 600xg and resuspended in cell-sorting medium [PBS, 25 mM HEPES pH7.0, 2% FBS (heat-inactivated) 1mM EDTA. 0.2 µm sterile-filtered]. Cell sorting was based on high-GFP-Alexa Flour 488 fluorescence; cells were sorted into a 96-well plate containing culture medium using a BD FACS Aria IIIu cell sorter.

At 10-14 days after sorting, the 96-well plates were examined for colonies. Wells with colonies were marked and allowed to grow for one week more before expanding. After 14 days, colonies were expanded from 96-well to 12-well plates using trypsin detachment. Cells were maintained in DMEM/F12 medium supplemented with 10% FBS and allowed to grow further. KO of VPS53 and VPS54 was confirmed by western blot analysis for absence of the targeted protein.

### Plasmid preparation, generation of lentiviral particles and stable cell lines

All constructs were generated using standard molecular biology techniques.

To construct mVps54-mCherry-Acta, DNA encoding mVPS54 was amplified by PCR using pCl-neo-mVPS54-13myc (24) as a template and following forward and reverse primers:

Forward: GCATCCGGAATGGCTTCAAGCCACAGTTC

Reverse: GGCGGTACCCCTCTTCTGTTCCCAGATCT

The amplified fragment was cleaved with BspE1 and KpnI and subcloned into mCherry-Acta (41) vector.

To generate pLenti COG4 Neo DEST construct, a chromosomal DNA fragment encoding the COG4 promoter region was amplified from human genomic DNA by PCR using the following forward and reverse primers:

Forward: GCTTATCGATTTCCCCCACGTCTGTTTACCA

Reverse: GAATTCTAGACTTGGTCCCCATTCGGCACTT

Then, the amplified fragment was cloned into pLenti CMV Neo DEST (705-1), using XbalI and ClaI as restriction sites. Lentiviral particles were generated using the HEK293FT cell line as described above.

To generate lentiviruses encoding GARP subunits, hVPS53 (GFP-tagged), transcript variant 1, (NM_001128159) purchased from Origene or mVps54-13myc (24) were subcloned into the pENTRA 1A no ccDB (w48-1) entry vector and recombined into pLenti COG4 promoter Neo DEST (705-1) destination vector using Gateway LR Clonase II Enzyme Mix (Thermo Fisher) according to the manufacturer’s instructions.

Lentiviral medium (100 µl) with polybrene (10 µg /ml) was used to transduce RPE1 cell lines in a six-well dish. At 24 h after transduction, the medium was replaced with 10% FBS without antibiotic in DMEM/F12. At 48 h after transduction, the medium was replaced with 600 µg/ml G418 selection medium and incubated for 72 h. Then, cells were single-sorted into a 96-well plate to obtain clonal populations. At 10-14 days after sorting, the 96-well plates were examined for colonies. Wells with colonies were marked and allowed to grow for one week more before expanding. Cells were maintained in DMEM/F12 medium with 10% FBS and allowed to grow further. Once colonies were split onto 10 cm dishes, aliquots were cryopreserved in freezing medium (90% FBS plus 10% DMSO).

### Preparation of cell lysates and western blot analysis

For preparation of cell lysates, cells grown on tissue culture dishes were washed twice with PBS and lysed in 2% SDS that was heated for 5 min at 70°C. Total protein concentration in the cell lysates was measured using BCA protein assay (Pierce). The protein samples were prepared in 6X SDS sample buffer containing beta-mercaptoethanol. The protein samples were denatured by incubation at 70°C for 10 minutes. 10-30 µg of protein samples were loaded into Bio-Rad (4–15%) gradient gel or Genescript (8–16%) gradient gel. Gels were then transferred onto nitrocellulose membrane using the Thermo Scientific Pierce G2 Fast Blotter. Membranes were rinsed in PBS, blocked in Odyssey blocking buffer (LI-COR) for 20 min, and incubated with primary antibodies overnight at 4°C. Membranes were washed with PBS and incubated with secondary fluorescently-tagged antibodies diluted in Odyssey blocking buffer for 60 min. Blots were then washed and imaged using the Odyssey Imaging System. Images were processed using the LI-COR Image Studio software.

### Lectin blotting

To perform blots with fluorescent lectins, membranes were blocked with 3% BSA, and stained with *Helix Pomatia* Agglutinin (HPA)-Alexa 647 (Thermo Fisher) or *Galanthus Nivalis* Lectin (GNL) conjugated to Alexa647 fluorophore (42) for 90 minutes and imaged using the Odyssey Imaging System.

### Secretion assay

Cells were plated in three 6-cm dishes and grown to 90-100% confluency. Cells were then rinsed 3 times with PBS and placed in 2 ml serum-free, chemically-defined medium (BioWhittaker Pro293a-CDM, Lonza) with 1× GlutaMAX (100× stock, Gibco) added per well. After 36 h, collected medium was spun down at 3,000xg to remove floating cells. Supernatant was concentrated using a 10k concentrator (Amicon® Ultra 10k, Millipore); final concentration was 24× that of cell lysates.

### Immunofluorescence assay and microscopy

Cells were plated on glass coverslips to 80-90% confluency and fixed with 4% paraformaldehyde (PFA) (freshly made from 16% stock solution) in phosphate-buffered saline (PBS) for 15 minutes at room temperature. Cells were then permeabilized with 0.1% Triton X-100 for one minute followed by treatment with 50 mM ammonium chloride for 5 minutes, treated with 6M urea for 2 minutes (only for COG3 staining) and washed with PBS. After washing and blocking twice with 1% BSA, 0.1% saponin in PBS for 10 minutes, cells were incubated with primary antibody (diluted in 1% cold fish gelatin, 0.1% saponin in PBS) for 40 minutes, washed, and incubated with fluorescently-conjugated secondary antibodies for 30 minutes. Cells were washed four times with PBS, then coverslips were dipped in PBS and water 10 times each, and mounted on glass microscope slides using Prolong® Gold antifade reagent (Life Technologies). Cells were imaged with a 63× oil 1.4 numerical aperture (NA) objective of a LSM880 Zeiss Laser inverted microscope and Airyscan super resolution microscope using ZEN software.

### RUSH assay

The RUSH construct Str-KDEL_flGalT-SBP-mCherry was transfected into cells using an electroporation-based system, and plated on glass coverslips in the presence of biotin-free media containing avidin (100 μg/ml) to prevent biotin in the medium from interfering with the RUSH reporter. At 16 h post-transfection, the medium was changed with the chase mix (biotin-free medium supplemented with biotin (40 μM) and cycloheximide (50 μM) for 2 h and 6 h, respectively, followed by fixing of cells with 4% PFA in PBS. Cells were stained for the ER maker PDI, Golgi marker GM130, endolysosomal marker CD63, and microscopic images were taken.

### Statistical analysis

All results are representative of at least 3 independent experiments. Western blot images are representative from 3 repeats. Western blots were quantified by densitometry using the LI-COR Image Studio software. Error bars for all graphs represent standard deviation. Statistical analysis was done using One-way ANOVA Graphpad Prism software.

## Results

### GARP KO in RPE1 and HEK293T cells causes N- and O-glycosylation defects

To test if complete depletion of GARP complex activity has an effect on the processing of N- and O-linked oligosaccharides of glycoproteins, we knocked out the VPS53 (Fig. 1A) and VPS54 (Fig. 1B) subunits of GARP in immortalized retinal pigment epithelial (RPE1) cells. These cells were chosen for their non-cancerous origin and superior characteristics for microscopy imaging. To test for protein N-glycosylation defects, non-permeabilized Wild Type (WT) and GARP-KO cells were stained with *Galanthus nivalis* (GNL) lectin labeled with Alexa647 (GNL647). GNL binds terminal 1,3 and 1,6 linked mannose residues on N-linked glycans (43) and an increase in binding to the plasma membrane indicates expression of underglycosylated N-linked glycoconjugates (10) (Fig. 1C). To determine O-glycosylation defects, non-permeabilized WT and GARP-KO cells were stained with *Helix pomatia* agglutinin (HPA) lectin labeled with Alexa647 (HPA647). HPA binds to terminal N-acetylgalactosaminyl residues in O-glycans; therefore, increased binding indicates expression of underglycosylated O-linked glycoconjugates (44) (Fig. 1D). GARP-KO cell lines showed a significant increase in binding of both lectins to the plasma membrane (Fig. 1C-F), indicating the presence of altered/immature N- and O-glycosylated proteins. To verify the specificity of the KO and of lectin staining, we rescued KO cells by stably expressing the corresponding GARP subunit under the control of the COG4 promoter. The use of the COG4 promoter (Zinia D’Souza and V.L, unpublished) allowed expression of COG and GARP subunits at near-endogenous levels (Fig. 1A and 1B). This expression level was sufficient to completely rescue the Cathepsin D sorting defect in GARP-KO cells (Fig. S1C). Importantly, in rescued cells, a substantial reduction in lectin binding to the plasma membrane was observed (Fig. 1C-F), confirming that glycosylation defects are directly related to GARP complex malfunction.

**Fig. 1.**
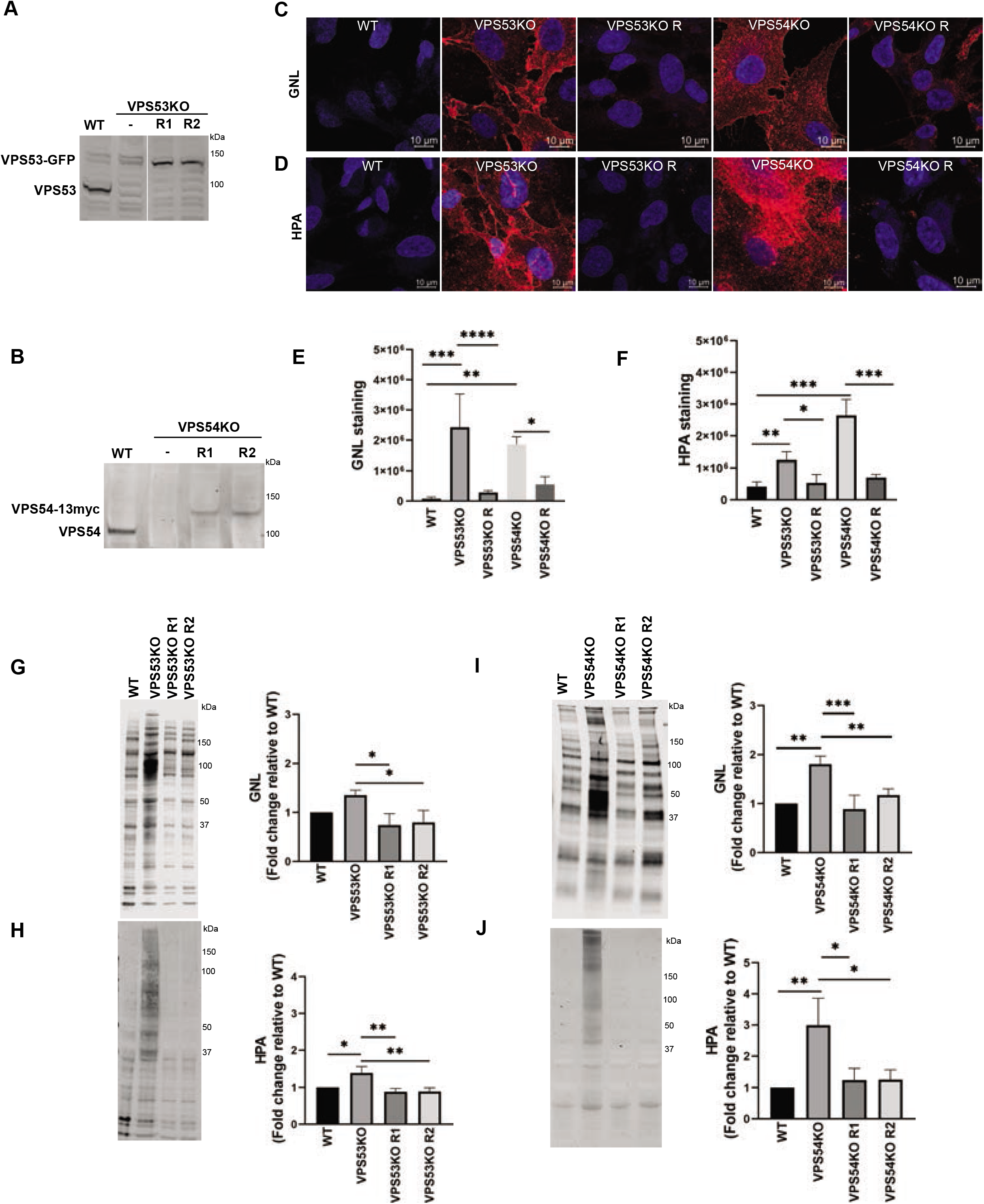
GARP-KO results in Golgi glycosylation defects. **A**, Western blotting (WB) of cell lysates from WT, VPS53-KO and two VPS53-KO rescued clones expressing VPS53-GFP (R1 and R2) probed with anti-VPS53 antibody. **B**, WB of cell lysates from WT, VPS54-KO and two VPS54-KO clones rescued with VPS54-13myc (R1 and R2) probed with anti-VPS54 antibodies. **C**, Staining of non-permeabilized WT, VPS53-KO and VPS54-KO cells, and the corresponding rescued cells with the fluorescently-conjugated lectin GNL-647 (specific for terminal α-D-mannosyl residues). **D**, Staining of non-permeabilized WT, VPS53-KO, VPS54-KO cells and the corresponding rescued cells with the fluorescently-conjugated lectin HPA-647 (specific for terminal GalNAc residues). **E**, Quantification of GNL-647 binding. Bar graphs with error bars represents mean ± SD from three different fields. **F**, Quantification of HPA-647 binding. Bar graphs with error bars represents mean ± SD from three different fields. **G, H**, GNL-647 and HPA-647 staining of total proteins from WT, VPS53-KO and two rescue clones (left panels), and quantification from three independent experiments (right panels). Values in bar graphs represent the mean ± SD from three independent experiments. Statistical significance was calculated in GraphPad Prism 8 using One-way ANOVA. **** P ≤ 0.0001, *** P ≤ 0.001, ** P ≤ 0.01, * P ≤ 0.05.

Lectin blot analysis using GNL and HPA also showed abnormal processing of N- and O-linked oligosaccharides in total cellular glycoproteins (Fig. 1G and 1J) and secreted glycoproteins (Fig. S1A and S1B) in VPS53- and VPS54-KO RPE1 cells. These glycosylation defects could also be rescued by stable expression of the corresponding genes in the KO cells (Fig. 1E-1J). From these experiments we concluded that KO of VPS53 or VPS54 in RPE1 cells caused N- and O-glycosylation defects in plasma membrane, intracellular and secreted glycoproteins. Similar glycosylation abnormalities were detected in VPS53- and VPS54-KO HEK293T cells (Fig. S1D and S1E), indicating that glycosylation defects in GARP-KO cells are cell line-independent and, hence, that the GARP complex plays a general role in oligosaccharide processing in human cells.

### GARP depletion affects the stability and glycosylation of Golgi and lysosomal glycoproteins

Defective glycosylation influences the stability of glycoproteins (42), so we reasoned that GARP deficiency might affect the stability of Golgi proteins. Indeed, we found that the levels of *cis*-Golgi GPP130 (Fig. 2A), medial-*trans*-Golgi TMEM165 (Fig. 2B) and TGN-resident TGN46 (Fig. 2C) were significantly reduced in GARP-KO cells. Importantly, the levels of all three Golgi proteins were restored in GARP-KO rescued cells. We also detected an electrophoretic mobility shift for all three proteins, indicating problems with their secondary modifications. The total level of the lysosomal protein LAMP2 (Fig. 2D) did not significantly change, but the mobility of LAMP2 was increased, indicating altered glycosylation of this protein. A similar pattern was observed in HEK293T cells, which also showed a decrease in the levels and increased electrophoretic mobility of GPP130 (Fig. S2A), TMEM165 (Fig. S2B) and TGN46 (Fig. S2C), and an increase in the mobility of LAMP2 (Fig. S2D). We concluded that the absence of GARP results in abnormal glycosylation of Golgi and lysosomal proteins, as well as reduced levels of Golgi proteins.

**Fig. 2.**
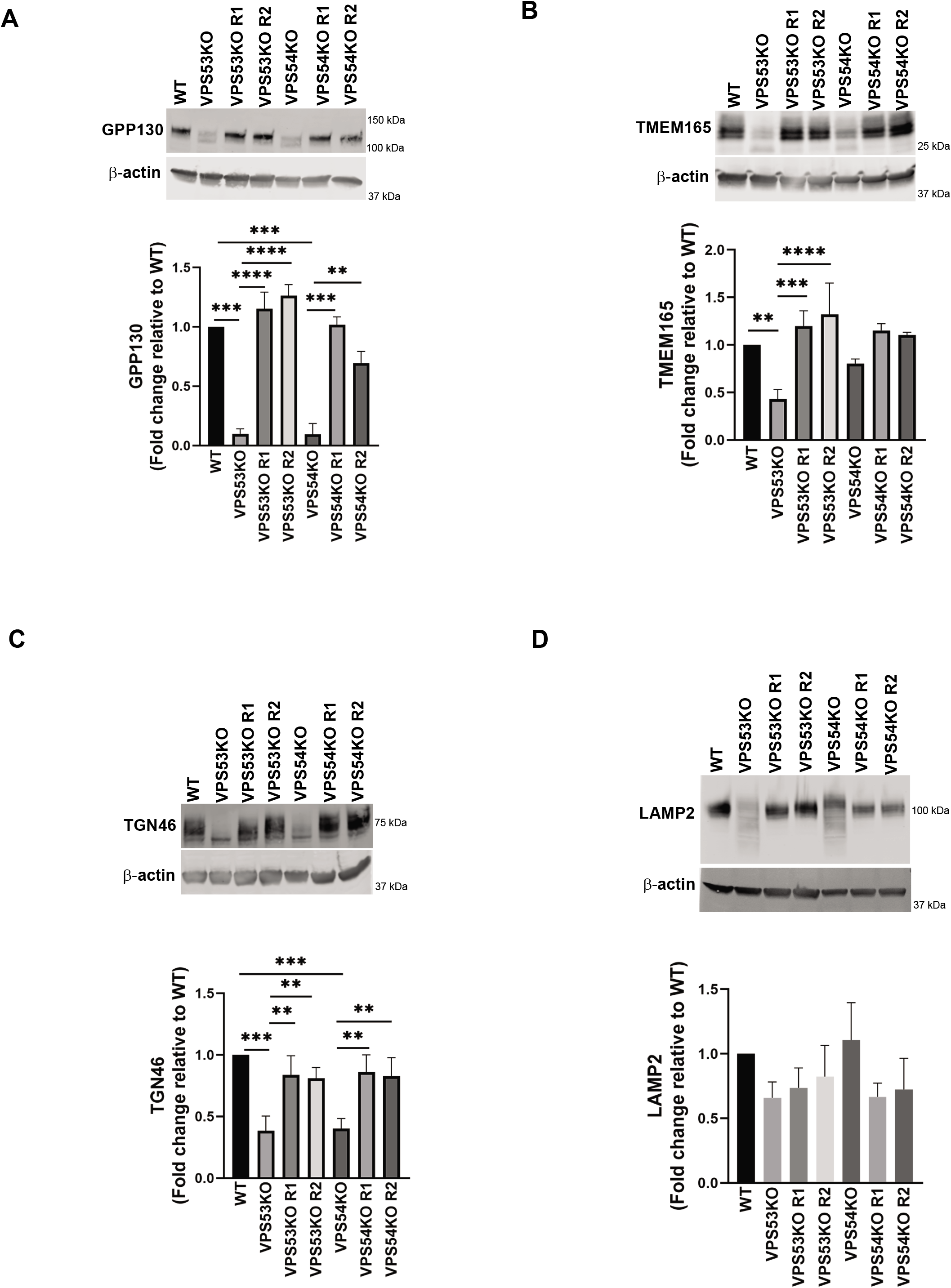
GARP-KO alters the glycosylation of Golgi and lysosomal glycoproteins in RPE1 cells. **A-D**, WB (top panel) and quantification (bottom panel) of GPP130 (**A**), TMEM165 (**B**), TGN46 (**C**) and LAMP2 (**D**) in WT, VPS53-KO, VPS54-KO and the corresponding rescued RPE1 cells. Quantification of the WB images mean ± SD was done by Image studio and graphs were prepared in GraphPad Prism 8. Values represent the mean ± SD from three independent experiments. Statistical significance was calculated using One-way ANOVA. **** P ≤ 0.0001, *** P ≤ 0.001, ** P ≤ 0.01.

### GARP KO affects the stability of key glycosylation enzymes in RPE1, HeLa and HEK293T cells

Defective Golgi glycosylation could result from malfunction, mislocalization and/or destabilization of Golgi enzymes. Indeed, VPS53- and VPS54-KO cells showed a significant reduction in the protein levels of the key N-glycosylation enzymes alpha-1,3-mannosyl-glycoprotein 2-beta-N-acetylglucosaminyltransferase (MGAT1) (Fig. 3A), beta-1,4-galactosyltransferase 1 (B4GalT1) (Fig. 3B) and beta-galactoside alpha-2,6-sialyltransferase 1 (ST6GalT1) (Fig. 3E). We also assessed the localization of B4GalT1 in WT, VPS53-KO and VPS53-KO rescued RPE1 cells by immunofluorescence microscopy (Fig. 3D left panel). We observed that the co-localization of B4GalT1 with the Golgi-marker GM130 was significantly decreased in VPS53-KO (Fig. 3D), indicating that GARP deficiency affects both the stability and localization of B4GalT1. We also examined polypeptide N-Acetylgalactosaminyltransferase 2 (GalNacT2), an O-glycosylation enzyme (45) and observed a decrease in the levels of this enzyme in VPS53- and VPS54-KO cells (Fig. 3C). Rescuing the VPS53-KO and VPS54-KO RPE1 cells by stable expression of the corresponding cDNAs significantly restored the levels and localization of N-and O-glycosylation Golgi enzymes (Fig. 3A-E).

**Fig. 3.**
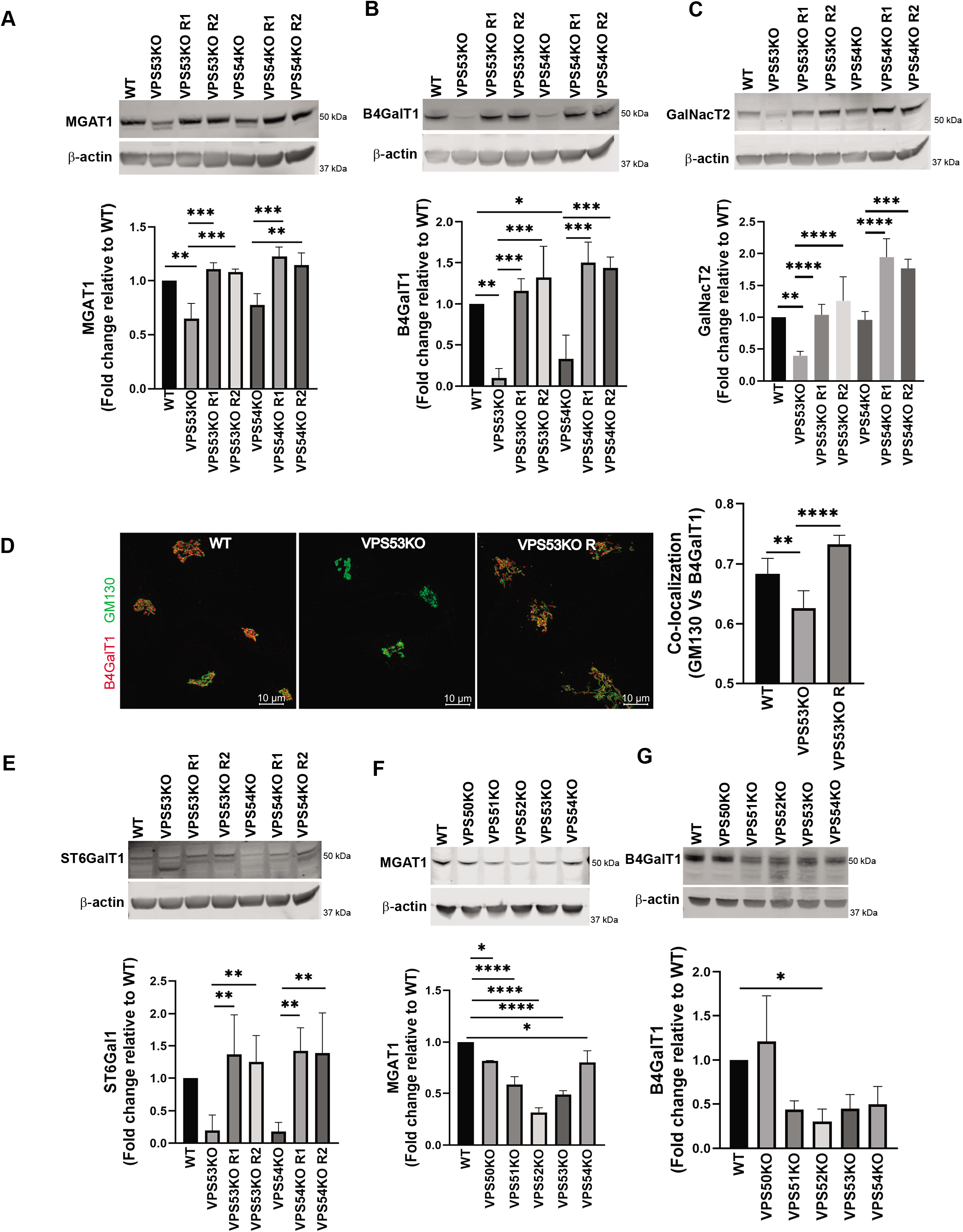
GARP-KO affects the stability of key N- and O- Golgi glycosylation enzymes in RPE1 and HeLa cells. **A-C**, WB (top panel) and quantification (bottom panels) of MGAT1 (**A**), B4GalT1 **(B**) and GalNAcT2 (**C**) in WT, VPS53-KO, VPS54-KO and the corresponding rescued RPE1 cells. **D**, WT, VPS53-KO and rescued RPE1 cells were co-stained for endogenous B4GalT1 (red) and GM130 (green), and images were taken (left panel). Co-localization of B4GalT1 with GM130 was determined by Pearson’s co-relation coefficient (right panel). At least 30 cells were imaged per sample for the quantification. **E**, WB (top panel) and quantification (bottom panels) of ST6Gal1 in WT, VPS53-KO, VPS54-KO and the corresponding rescued RPE1 cells. **F**,**G**, WB (top panel) and quantification (bottom panels) of MGAT1 (**F**) and B4GalT1 (**G**) in WT, VPS50-, VPS51-, VPS52-, VPS53- and VPS54-KO HeLa cells. Values in bar graphs represent the mean ± SD from three independent experiments. Statistical significance was calculated using One-way ANOVA. **** P ≤ 0.0001, *** P ≤ 0.001, ** P ≤ 0.01, * P ≤ 0.05.

In line with these findings, KO of any of the GARP subunits (VPS51, VPS52, VPS53 and VPS54) in HeLa cells (24) also caused a significant reduction of MGAT1 (Fig. 3F) and B4GalT1 protein levels (Fig. 3G). In contrast, KO of the VPS50 subunit of EARP had little or no effect on the protein levels of MGAT1 (Fig. 3F) and B4GalT1 (Fig. 3G), indicating that glycosylation defects are due to malfunction of the GARP and not EARP complex. We also observed reduced proteins levels of MGAT1 (Fig. S3A) and B4GalT1 (Fig. S3B) in VP53- and VPS54-KO HEK293T cells. In contrast, the protein levels of ST6Gal1 were not reduced, but its electrophoretic motility was altered, in VP53- and VPS54-KO HEK293T consistent with defects in ST6Gal1 modification and/or with partial degradation.

The defects in Golgi enzyme stability observed in GARP-KO cells are very similar to those previously described in COG-KO cells (12, 42), so we tested if GARP depletion affects COG localization and/or stability. We found that this was not the case, as both localization (Figure S4A) and stability (Figure S4B) of COG subunits were unaltered in GARP-KO cells. We concluded that depletion of the GARP complex specifically affects the stability of key glycosylation enzymes in all tested human cell lines.

### ST6Gal1 is not retained in the Golgi in GARP-KO cells

The GARP complex is localized to the TGN (7); therefore, we hypothesized that in GARP-deficient cells newly-synthesized glycosylation enzymes may not be efficiently retained in and/or recycled to the Golgi cisternae and may instead be targeted to endolysosomes. To test this hypothesis, we transfected WT and VPS53- and VPS54-KO RPE1 cells with plasmids encoding RFP-tagged ST6Gal1 (ST-RFP) and GFP-tagged LAMP2 (LAMP2-GFP), and examined the localization of both proteins relative to the Golgi marker GM130 at 4 and 20 hours after transfection. As expected, 4 hours were sufficient for ST-RFP to become localized to the Golgi complex in both WT and GARP-KO cells (Fig. 4A). At 20 hours after transfection, most of the ST-RFP remained localized to the *trans*-Golgi in WT cells (Fig. 4B). In contrast, in VPS53- and VPS54-KO cells, ST-RFP exhibited partial localization to cytoplasmic puncta, the majority of which co-localized with LAMP2-GFP (Fig. 4B). This result revealed that ST-RFP is not efficiently retained in Golgi in GARP-KO cells but is instead targeted to the endolysosomal compartment for degradation.

**Fig. 4.**
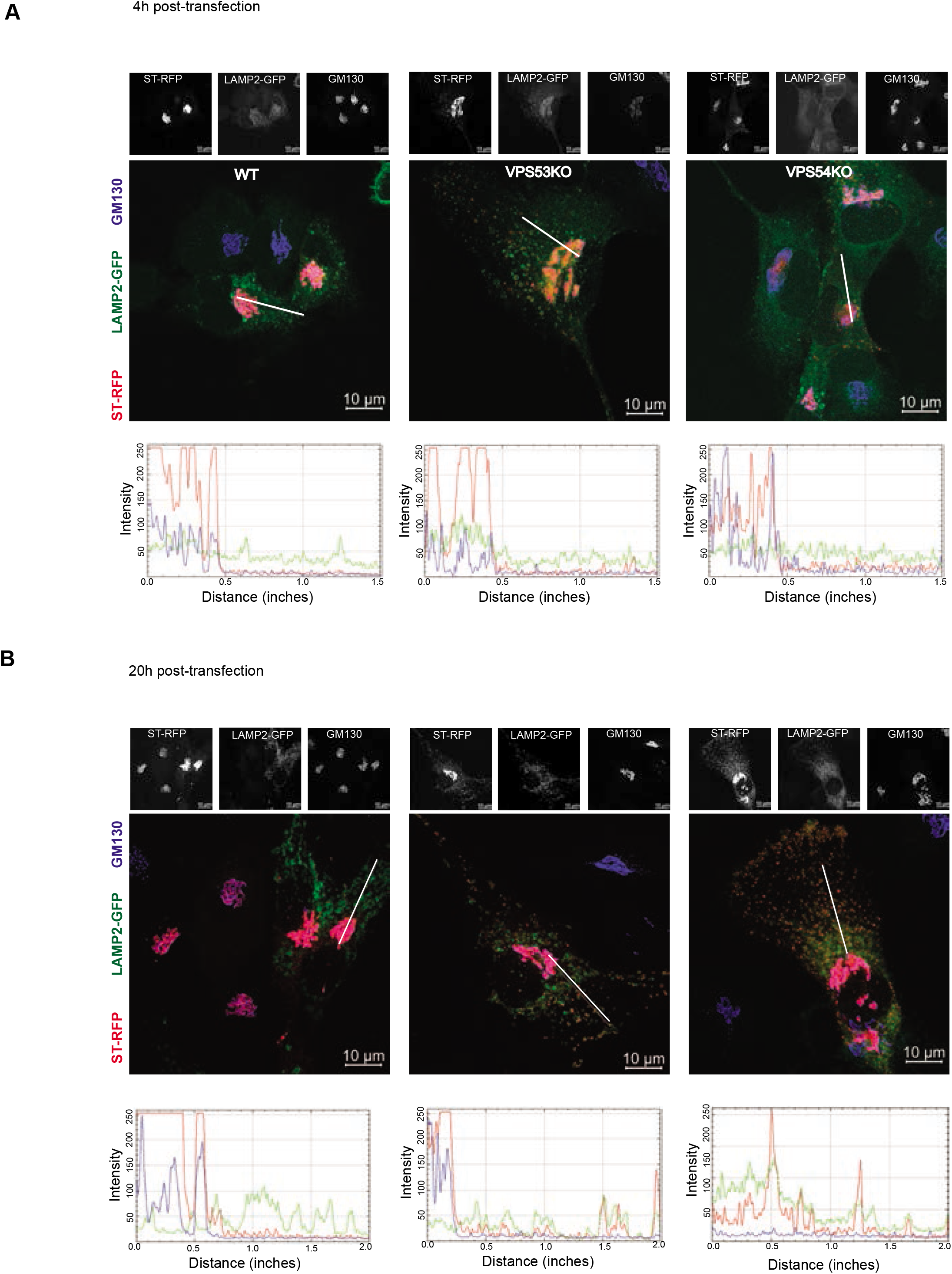
ST-RFP is not retained in the Golgi in GARP deficient cells. **A**,**B**, WT, VPS53-KO, and VPS54-KO RPE1 cells were transfected with plasmids encoding ST-RFP and LAMP2-GFP for 4h (A) and 20h (**B**), followed by staining for the Golgi marker GM130. Microscopic images for both **A**, and **B** show individual channels in grayscale on the top of each merged image. Line scan plots of relative intensity over distance show overlap between the channels (bottom panel).

### RUSH assay reveals fast mislocalization of B4GalT1 in GARP-KO cells

We next employed the Retention Using Selective Hooks (RUSH) system (40) to examine the transport characteristics of the Golgi enzyme B4GalT1 in WT and GARP-KO cells. RPE1 cells were transfected to express streptavidin-KDEL as a hook and B4GalT1-SBP-mCherry as a reporter. Following overnight transfection, biotin was added to release GalT-SBP-mCherry from ER retention and samples were analyzed at 0, 2 and 6 hours to compare the intracellular trafficking of B4GalT1 in WT and GARP-KO cells. Staining for the ER marker PDI (shown in green) revealed, as expected, that B4GalT1 (shown in red) was localized to the ER in the absence of biotin (0 hours) in WT, VPS53-KO and VPS54-KO cells (Fig. 5A). Two hours after biotin addition, B4GalT1 moved to the perinuclear area, where it co-localized with the *cis*-Golgi marker GM130 (Fig. 5B) in both WT and GARP-KO cells, indicating that ER-Golgi traffic is unaltered in these cells. Six hours after release from the ER (Fig. 5C, left panel), B4GalT1 was localized to the Golgi area in WT cells. In contrast, VPS53-KO and VPS54-KO cells showed a dramatic decrease in B4GalT1 co-localization with GM130 (Fig. 5C, right panel) and the appearance of B4GalT1 in multiple puncta that partially co-localized with the endolysosomal marker CD63 (Fig. 5D). Similar results were obtained by performing the RUSH assay in HeLa cells (Figure S5). In this case, both B4GalT1 (Fig. S5 right panel) and MAN2A (Fig. S5 left panel) became partially mislocalized to endolysosomes in VPS54-KO cells. To further test that mislocalization of B4GalT1 is due to GARP defect, we have employed mitochondria mislocalization strategy, that was previously used to re-localize Golgi-derived vesicles to mitochondria decorated with COG4/COG8 tagged with mitochondria targeted Acta sequence (41). VPS54-mCherry was C-terminally tagged with the mitochondria targeting sequence – (VPS54-mCherry-Acta) and introduced in WT and VPS54-KO cells (Fig. S6A). In both cell types, VPS54-mCherry-Acta was expressed in a typical mitochondria network pattern. Staining cells with antibodies to B4GalT1 revealed that in both untransfected cells and in WT cells transfected with VPS54-mCherry-Acta B4GalT1 was mostly localized to the Golgi, while in VPS54-KO cells this enzyme showed a more dispersed pattern of localization with an increased co-localization with VPS54-mCherry-Acta. This result suggests that the GARP complex may directly tether transport intermediates that recycle Golgi enzymes. The nature of these trafficking intermediates and the exact mechanism of GARP-dependent trafficking of Golgi enzymes will be a subject of our future investigation.

**Fig. 5.**
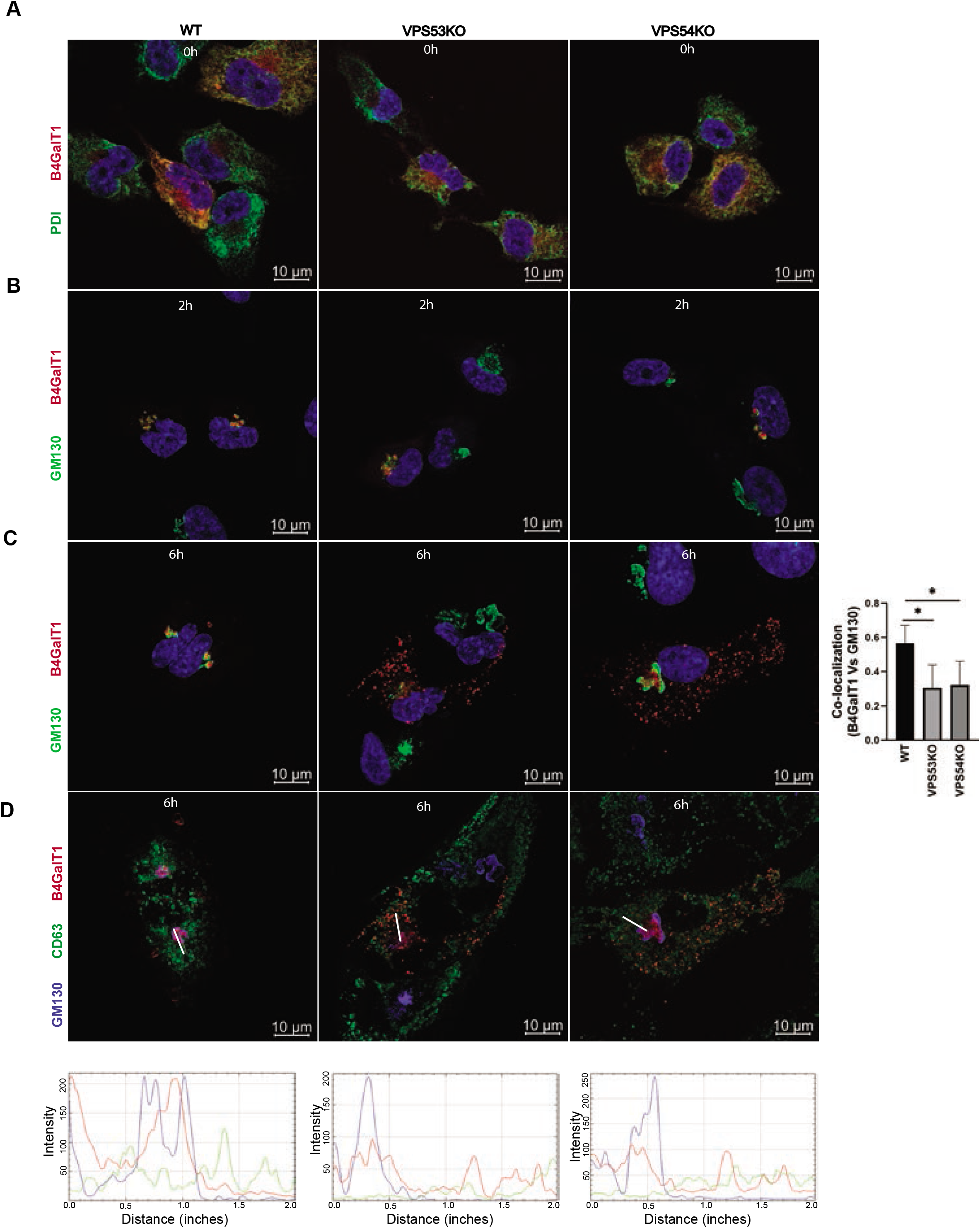
RUSH assay reveals mislocalization of B4GalT1 to endolysosomes in GARP-KO cells. RPE1 cells were transfected with plasmids encoding Str-KDEL_flB4GalT1-SBP-mCherry and chased with biotin mix for 0h, 2h, and 6h. **A**, Co-localization of B4GalT1 with ER marker PDI at 0h. **B**,**C**, Co-localization of B4GalT1 with Golgi protein GM130 at 2h (**B**) and 6h (**C**). The graph on the right side of panel **C** shows the quantification of B4GalT1 and GM130 co-localization at 6h of biotin chase. Values in the bar graph represent the mean ± SD from the co-localization between B4GalT1 and GM130 from 30 different cells. Statistical significance was calculated using One-way ANOVA. * P ≤ 0.05. **D**, Co-localization of B4GalT1, GM130 and the endolysosomal marker CD63 after 6h of chase. The bottom panels are line scan plots of relative intensities against distance for all the channels.

Taken together, these results indicate that GARP is essential for proper localization of Golgi enzymes in different cell types.

## Discussion

In this study, we have demonstrated that the GARP complex is important for the proper modification of oligosaccharide chains of glycoproteins in the Golgi complex. Using multiple cell lines, we discovered that complete depletion of two GARP subunits (VPS53 and VPS54) is detrimental for the modification of both N- and O-linked oligosaccharides and the stability of glycoproteins and components of the Golgi glycosylation machinery.

In addition to the Golgi glycosylation defect, our results also demonstrated reduction in the levels of Golgi glycoproteins and increase in electrophoretic mobility of LAMP2 in GARP-KO cells. We also observed alterations in the localization and stability of multiple Golgi glycosyltransferases in GARP-KO cells. We propose that glycosylation defects in GARP-KO cells are directly linked to altered stability of the glycosylation machinery due to mislocalization of Golgi glycosyltransferases. The stability of Golgi enzymes was unaltered in VPS50-KO cells, indicating that only GARP (VPS51-54), and not EARP (VPS50-53), is needed for the maintenance of the Golgi glycosylation machinery. There might be several reasons behind the reduction in stability of Golgi resident glycosyltransferases in GARP-KO cells. One reason might be a failure in the retrieval of glycosyltransferases to their corresponding Golgi cisternae. As a consequence, these Golgi enzymes would be unable to maintain their proper steady-state localization and thus meet their substrates. This would be similar to COG subunit mutants, which exhibit a reduction in the Golgi localization of B4GalT1 and MGAT1 (42) (46). Another reason might be the mislocalization of Golgi resident enzymes away from the Golgi to other compartments. This would be similar to observations in COG4-KO cells, which showed mislocalization of the Golgi enzyme ST6Gal1 tagged with RFP to Enlarged Endolysosomes (EELs), indicating that these Golgi resident enzymes can be degraded in EELs (12). Our results showed increased colocalization of ST6Gal1 with LAMP2 (Figure 4B) and B4GalT1 with CD63 (Figure 5D), in agreement with the COG-KO studies.

Glycosylation abnormalities were previously described in cells depleted for another Golgi vesicle tethering complex, COG (10), and GARP and COG were shown to functionally interact via VPS51 (47) and a STX16-containing SNARE complex (48) (49). Therefore, one possibility was that GARP depletion affects COG complex function.

However, both the localization and stability of COG complex subunits were unaltered in GARP-KO cells, indicating that the glycosylation defects observed in this study are directly related to GARP. Side-by-side comparison of glycosylation defects in HEK293T and RPE1 cells depleted for multiple COG and GARP subunits showed that stability of the tested Golgi enzymes was altered to a comparable level, but the N- and O-glycosylation defects were less severe in GARP-KO cells (A.K., T.K, V.L., unpublished), indicating that COG deficiency affects a wider range of Golgi glycosylation machinery.

The effect of GARP depletion on the Golgi glycosylation machinery was unexpected, since current models for trafficking and retention of Golgi enzymes include only intra-Golgi and Golgi-ER cycling pathways (50) (51) (18). Golgi-resident proteins are concentrated in “proper” Golgi sub-compartments by a combination of retention and retrieval mechanisms (52), but for the majority of Golgi enzymes the localization mechanisms are still enigmatic. One model proposed that the thickness of the lipid bilayer is important for the proper localization and retention of transmembrane proteins (53). Other models rely on efficient intra-Golgi and Golgi-ER recycling (54) (55). Recycling is achieved by packaging of Golgi resident proteins into recycling COPI vesicles (56), although COPI-independent Golgi-ER recycling was also described (57). A small subset of enzymes depends on direct interaction with COPI (58), while another subset uses the GOLPH3 adaptor for interaction with COPI (59). There are multiple scenarios for how GARP deficiency may affect the maintenance of Golgi enzymes. In the “indirect” models, GARP deficiency may affect the thickness of TGN membranes (by somehow altering lipid balance in this compartment) or proper retrieval of enzyme adaptors. Our preliminary analysis indicates that localization of COPI coat was not severely altered in GARP-KO cells and that glycosylation defects observed in GOLPH3/GOLPH3L double-KO cells are less severe than in GARP-depleted cells (T.K., V.L. unpublished observation). It is important to note that GARP deficiency affects not only enzymes located at the *trans*-Golgi (B4GalT1 and ST6Gal1), but also enzymes located in earlier Golgi compartments (MGAT1 and GalNacT2). Although both MGAT1 and B4GalT1 were found in COPI vesicles, these enzymes do not bind COPI coat directly and the nature of a possible adaptor is still unknown. Furthermore, a number of factors, including pH and ion concentration of the lumen of each cisterna, contribute to the fidelity of glycosylation. The recycling of Golgi-resident ion channels and transporters may also be affected by the loss of GARP function, leading to overall disruption of Golgi homeostasis.

Interestingly, live-cell imaging of WT HeLa cells stably expressing B4GalT1-GFP and ST-RFP revealed that both enzymes appeared not only in the Golgi cisternae and multiple vesicular structures, but also in the tubular sorting/late endosome-like compartment (Movie S1), indicating that this compartment could be a part of the normal trafficking itinerary for Golgi enzymes, and that defects in GARP-mediated retrieval from this compartment may cause mislocalization and degradation of the enzymes, causing numerous glycosylation defects (Fig. 6). If this hypothesis is correct, we would expect that the GARP complex may interact with vesicles that recycle Golgi enzymes. Indeed, relocalization of VPS54 to mitochondria in VPS54-KO cells caused dispersal of B4GalT1 signal with partial co-localization of B4GalT1 vesicles with Vps54 decorated membranes.

**Fig. 6.**
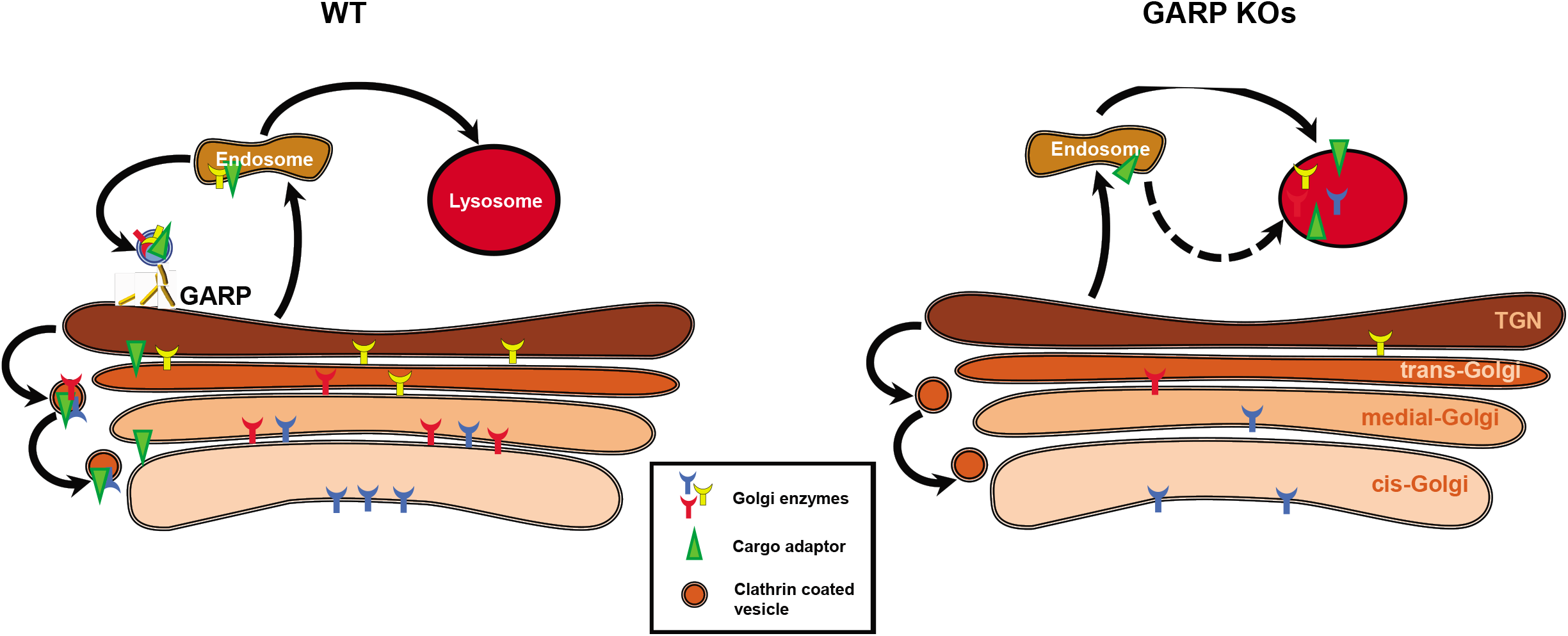
Proposed model of altered Golgi trafficking in GARP-KO cells. GARP complex plays a key role in recycling of Golgi enzymes from endosome to the TGN. In WT cells (left), the Golgi enzymes that modify newly-synthesized glycoproteins are recycled back to their corresponding Golgi compartments by GARP, allowing them to maintain their proper steady-state localization. In contrast, in GARP-KO cells (right) Golgi enzymes are not recycled back and are instead delivered to lysosomes for degradation. Consequently, there are decreased levels of enzymes in the Golgi and carbohydrate modification of glycoproteins is hindered.

In summary, we have demonstrated that the GARP-KO human cells have defects in protein N- and O-glycosylation. Moreover, GARP-KO cells are unable to retain the tested Golgi glycosylation enzymes. Hence, the GARP complex is a new component of the cellular machinery that regulates the proper localization and abundance of Golgi enzymes.

## Abbreviations used

B4GalT1: Beta-1,4-galactosyltransferase 1
COG: Conserved Oligomeric Golgi
GARP: Golgi-associated retrograde protein
GalNacT2: N-acetyl galactosaminyltransferase 2
MTC: Multi-subunit Tethering Complex
MGAT1: Alpha- 1,3-Mannosyl-Glycoprotein 2-Beta-N-Acetylglucosaminyltransferase
RUSH: Retention Using Selective Hooks
SNARE: SNAP (Soluble NSF Attachment Protein) Receptor
ST6Gal1: Beta-galactoside alpha-2,6-sialyltransferase 1
TGN: trans-Golgi Network
VPS: Vacuolar Protein Sorting

## Acknowledgments

We are thankful to Eric Campeau, Rainer Duden, Wei Guo, Paul Kaufman, Taroh Kinoshita, James Rothman, Frank Perez, Santiago M. Di Pietro, Didier Trono and others who provided reagents and cell lines. We would also like to thank the UAMS sequencing, flow cytometry and digital microscopy core facilities for the use of their equipment and expertise. We are grateful to all members of Lupashin’s lab for comments on the manuscript. This work was supported by the National Institute of Health (R01GM083144) and by UAMS Easy Win Early Victory grant program (VL), and by the Intramural Program of NICHD (ZIA HD001607).

## Autor contributions

A.K., T.K. and V.L. devised and planned the study. A.K. and T.K. performed the experiments. A.K.,T.K., J.B. and V.L. wrote and edit the manuscript.

## Supplementary figure legends

**Supplementary Fig. S1. GARP KO alters glycosylation in secreted glycoproteins and total lysates. A**, GNL-647 staining of secretory proteins from WT, GARP-KO and rescued RPE1 cells. **B**, HPA-647 staining of secretory proteins from WT, GARP-KOs and rescued RPE1 cells. **C**, WB of cathepsin D secreted from WT, GARP-KO and rescued RPE1 cells. Representative images from three independent experiments are shown. **D**,**E**, GNL-647 (**D**) and HPA-647 (**E**) staining of whole-cell lysates from WT, GARP-KO HEK293T cells (left) and quantification of staining relative to WT HEK293T cells (right). Bars represent the mean ± SD of values from three independent experiments. Statistical significance was calculated using One-way ANOVA. ** P ≤ 0.01, *** P ≤ 0.001.

**Supplementary Fig. S2. GARP KO alters the glycosylation of Golgi and lysosomal glycoproteins in HEK293T cells. A-D**, WB of endogenous GPP130 (**A**), TMEM165 (**B**), TGN46 (**C**) and LAMP2 (**D**) from cell lysates of WT, VPS53-KO and VPS54-KO cells.

**Supplementary Fig. S3. GARP KO alters the stability of N- and O-Golgi glycosylation enzymes in HEK293T cells. A-C**, WB (top) and quantification (bottom) of MGAT1 (**A**), B4GalT1 (**B**) and ST6Gal1 (**C**). Bars represent the mean ± SD of values from three independent experiments. Statistical significance was calculated using One-way ANOVA. **** P ≤ 0.0001, ** P ≤ 0.01.

**Supplementary Fig. S4. Localization of COG3 or COG8 is not altered in GARP-KO cells**. (**A**) RPE1 cells were stained for COG3 (top row) or COG8 (bottom row) together with GM130, and images were taken by super resolution Airyscan microscopy. (**B**) WB of COG3 and COG8 in GARP-KO and rescued cells.

**Supplementary Fig. S5. RUSH assay reveals mislocalization of B4GalT1 in GARP-KO HeLa cells**. WT and VPS54-KO HeLa cells were transfected with plasmids encoding a B4GalT1 RUSH construct and chased with biotin mix for 1h (left panels) and 6h (right panels), followed by staining for MAN2A and Giantin. The arrows on the right panel indicate the co-localization of B4GalT1 and MAN2A.

**Supplementary Fig. S6. Relocation of VPS54 subunit of GARP complex to mitochondria impairs the Golgi enzyme B4GalT1 localization in VPS54 KO RPE1 cells**. (**A**) WT and VPS54KO RPE1 cells were transfected with VPS54 mCherry Acta and stained for B4GalT1. On the right side, the inverted image of B4GalT1 individual channel in WT and VPS54KO RPE1 cells are shown. (**B**) Quantification of 20 transfected cells for co-localization between VPS54-mCherry-Acta and B4GalT1 were used in both WT and VPS54KO samples using GraphPad prism. **** P ≤ 0.0001.

