## Supplemental figures 1-6 for "The Golgi-associated retrograde protein (GARP) complex plays an essential role in the maintenance of the Golgi glycosylation machinery"

### Supplementary 1

S1A

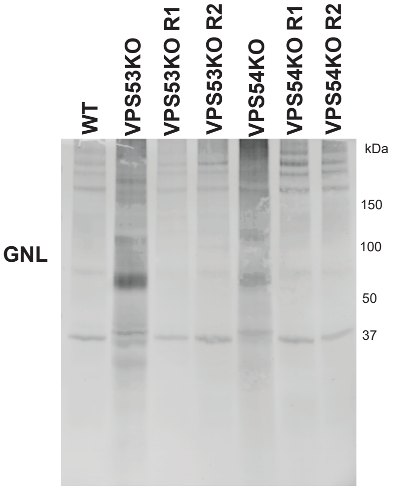


S1B

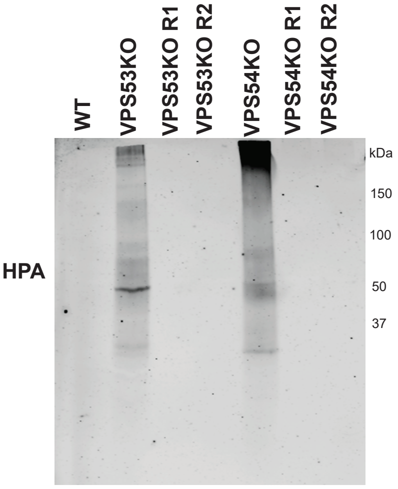


S1C

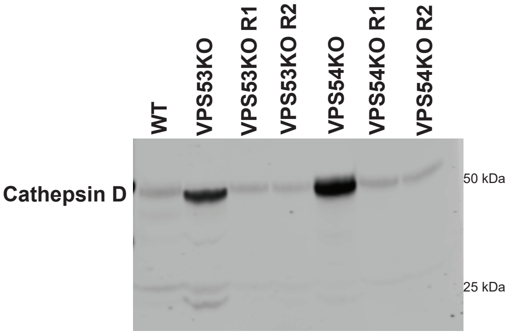


S1D

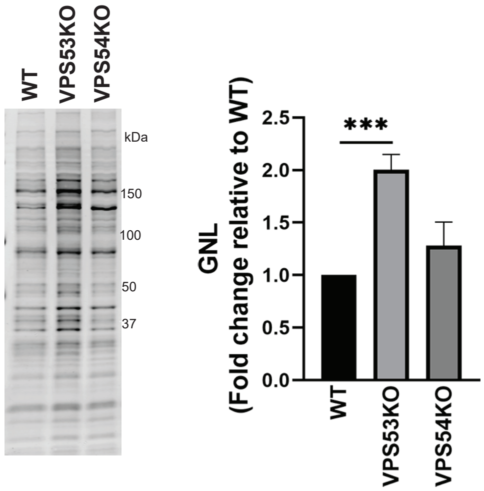


S1E

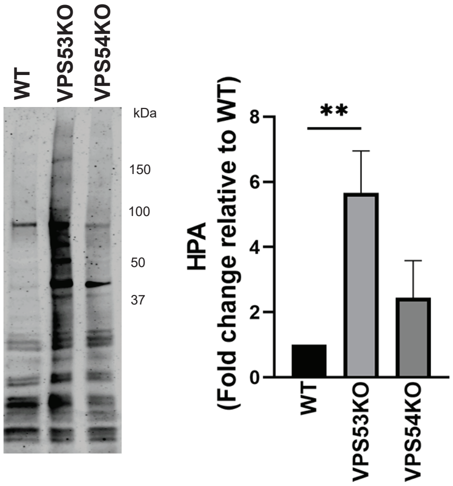

#### Supplementary 2

### S2A

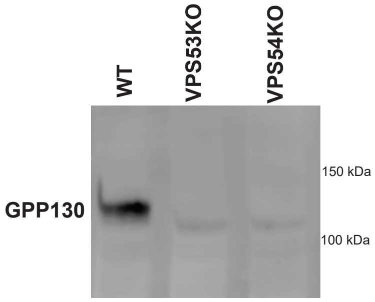


### S2B

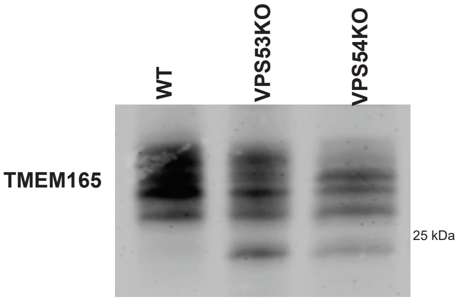


### S2C

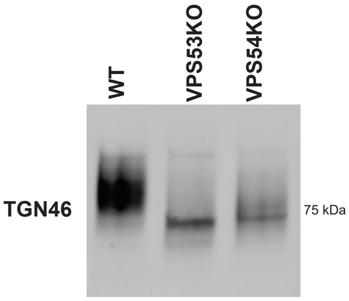


### S2D

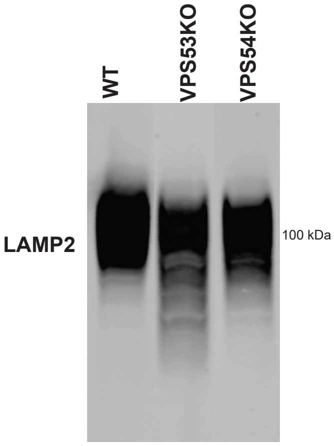

**Supplementary 3**

## S3A

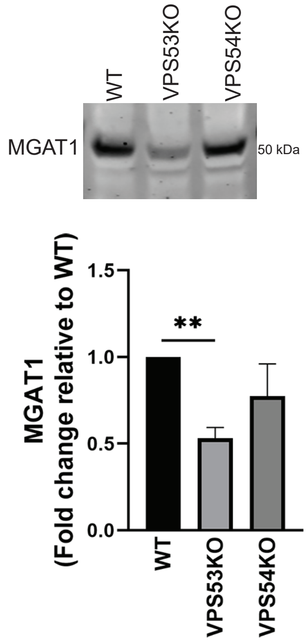


## S3B

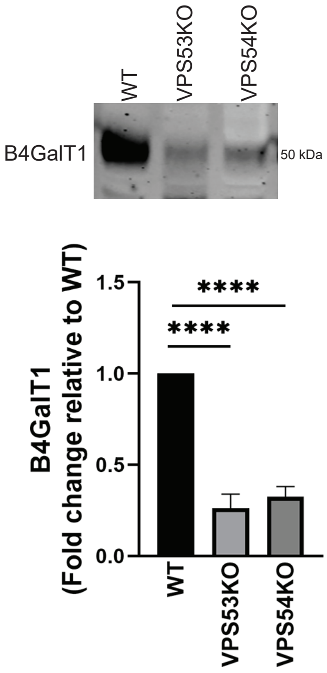


## S3C

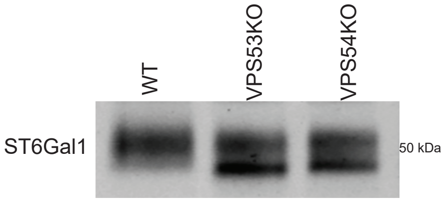

#### Supplementary 4

### S4A

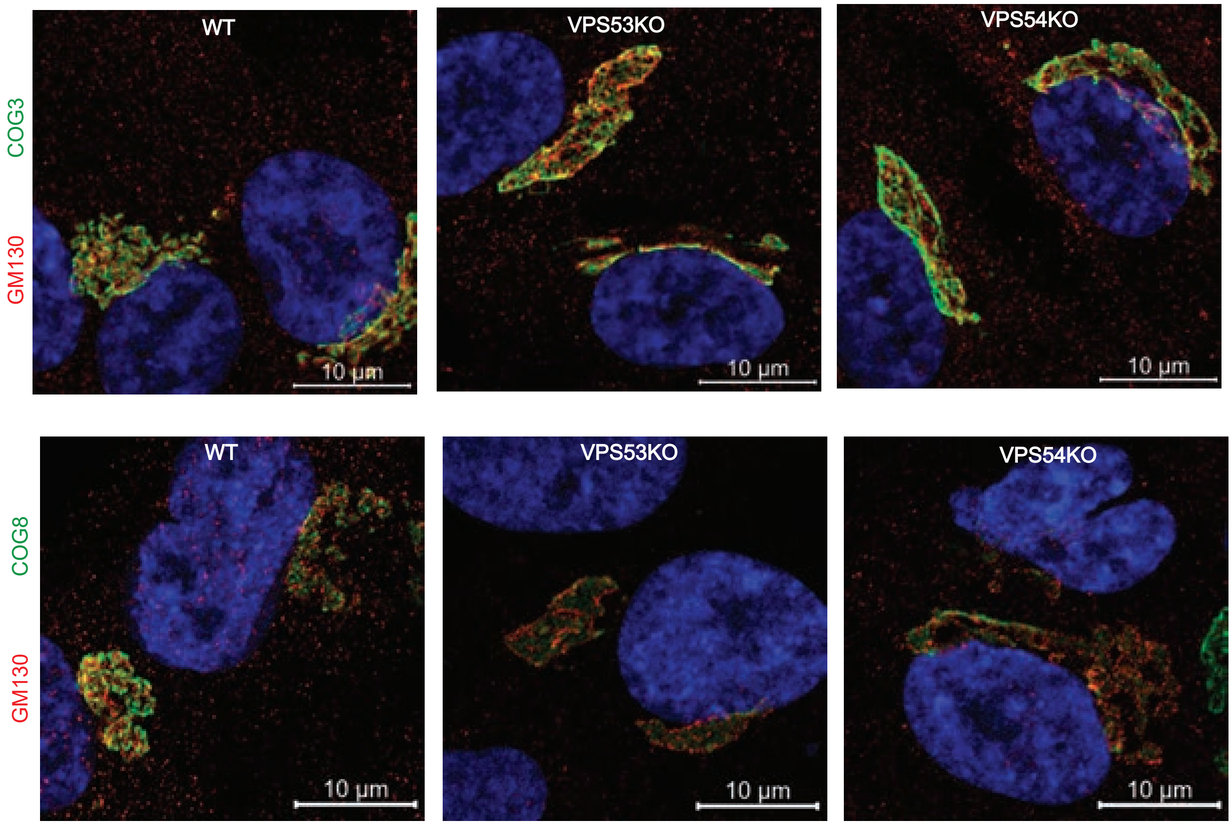


### S4B

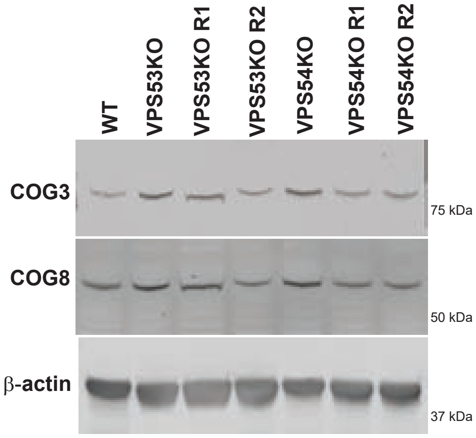

### Supplementary 5

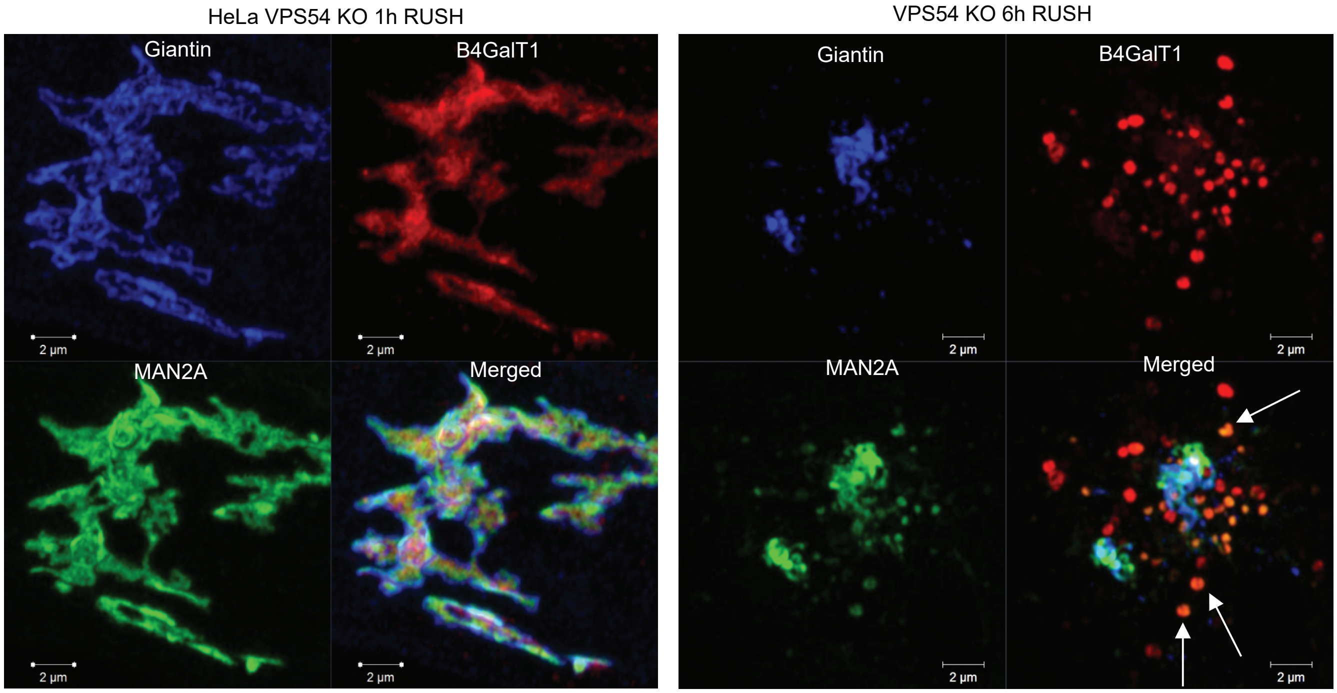

#### Supplementary 6

### S6A

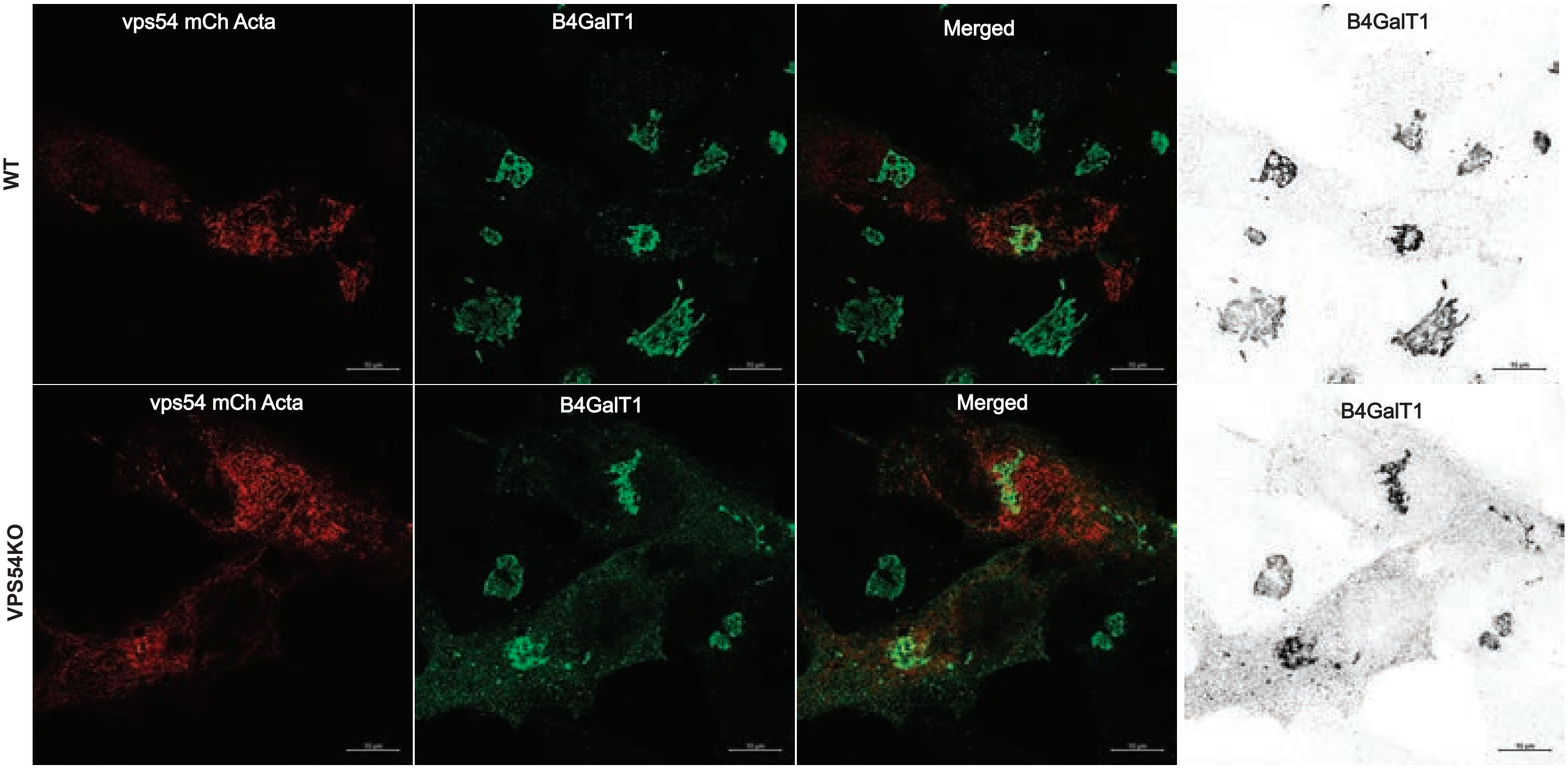


### S6B

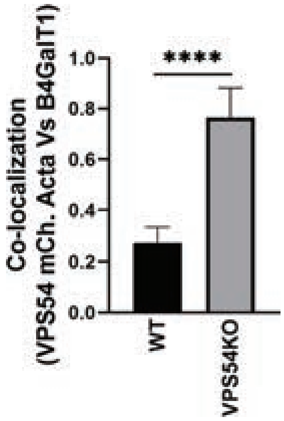
